# Degradation of miR-466d-3p by JEV NS3 facilitates viral replication and IL-1β expression

**DOI:** 10.1101/681569

**Authors:** Hui Jiang, Caiquan Zhao, Zhuofang Bai, Yanqing Meng, Tian Qin, Xiao Wang, Guojun Wang, Min Cui, Jing Ye, Shengbo Cao, Guangpeng Li, Yang Yang

## Abstract

Previous studies revealed that Japanese encephalitis virus (JEV) infection alters the expression of miRNA in central nervous system (CNS). However, the mechanism of JEV infection contributes to the regulation of miRNAs in CNS remain obscure. Here, we found that a global degradation of mature miRNA in mouse brain and neuroblastoma cells after JEV infection. In additional, the integrative analysis of miRNAs and mRNAs suggests that those down-regulated miRNAs are primarily targeted inflammation genes and the miR-466d-3p target the IL-1β which in the middle of those inflammation genes. Transfection of miR-466d-3p decreased the IL-1β expression and inhibited the JEV replication in NA cells. Interestingly, the miR-466d-3p level increased after JEV infection in the presence of cycloheximide, which indicated that viral protein expression reduces miR-466d-3p. Therefore, we generated all the JEV coding protein and demonstrated that NS3 is a potent miRNA suppressor. Furthermore, the NS3 of ZIKA virus, WNV, DENV1 and DENV2 also decreased the expression of miR-466d-3p. The in vitro unwinding assay demonstrated that the NS3 could unwind the pre-miR-466d and induce the disfunction of miRNA. Using computational models and RNA immunoprecipitation assay, we found that arginine-rich domains of NS3 are critical for pre-miRNA binding and the degradation of host miRNAs. Importantly, site-directed mutagenesis of conserved residues revealed that R226G and R202W significantly reduced the binding affinity and degradation of pre-miR-466d. Together, these results extend the helicase of Flavivirus function beyond unwinding duplex RNA to the decay of pre-miRNAs, which provides a new mechanism of NS3 in regulating miRNA pathways and promoting the neuroinflammation.

**Author Summary:** Host miRNAs had been reported to regulate JEV induced inflammation in central nervous system. We found that the NS3 of JEV can reduce most of host miRNA expression. The helicase region of the NS3 specifically binds to precursors of miRNA and lead to incorrect unwinding of precursors of miRNAs which inhibits the function of miRNAs. This observation leads to two major findings. First, we identified the miR-466d-3p targets to the host IL-1β and E protein of JEV, and NS3 degrades the miR-466d-3p to promote the brain inflammation and viral replication. Second, we proved that the arginine on the helicase of NS3 is the main miRNA binding sites, and the miRNA degradation by NS3 was abolished when the R226 and R202 were mutated on the NS3. These findings were also confirmed with NS3 of ZIKA virus, WNV and DENV which could decrease the expression level of miR-466d-3p to enhance the inflammation. Our study provides new insights into the molecular mechanism of encephalitis caused by JEV, and reveals several amino acid sites to further attenuate the JEV vaccine.

## Introduction

JEV is a single-stranded, positive-sense RNA virus belonging to flavivirus of the Flaviviridae family. Its genome encodes three structural proteins in order of envelope (E), capsid (C), and premembrane (PrM), and seven nonstructural (NS) proteins, NS1, NS2A, NS2B, NS3, NS4A, NS4B, and NS5. During the JEV life cycle in cells, the replication was initiated with the RNA genome replicase complex, of which NS3 and NS5 are major components required for genome replication and capping (11). The polyprotein NS3 of JEV belongs to superfamily 2 helicase, which has protease, nucleoside triphosphatase, ATP-dependent RNA helicase activities and unwind double strands genome during the viral replication (63).

JEV is a neurotropic virus causing neuroinflammation and neuronal damage that causes Japanese encephalitis (JE) (44). In spite of vaccines available control JEV, approximately 67,900 cases of JE are reported around the world per year, half of which were occurred in the mainland of China (7). Due to the lack of effective anti-viral drug against JEV, the fatality rate of JE is ∼25-32%, and 50% of the survivors suffer from neuropsychiatric sequelae (55). JEV causes neuroinflammation and neuronal damage in mammalian hosts by modulating cytokine/chemokine production (10) as well as the activation and migration of neuroglial cells (12, 19). The production of inflammation cytokine increases microglial activation following JEV infection, which facilitates outcome of viral pathogenesis and benefit the dissemination of the virus in the CNS (25, 35). For this reason, most of studies on innate immune response in the CNS have focused on the molecular components of JEV recognition and signaling regulator. Recent studies have revealed that the JEV was recognized by the toll-like receptor 7 (TLR7) in neuron cells and TLR3 in microglia cells (28, 47). Following JEV recognition, the adapter proteins of TLRs are activated and involved myeloid differentiation primary response 88 (MyD88). Furthermore, protein kinases such as kinase B (Akt), phosphoinositide 3-kinase (PI3K), p38 mitogen-activated protein kinase (p38MAPK) and signal-regulated kinase (ERK) (18) are triggered by JEV infection. However, the involvement of JEV components in modulating inflammation response still obscure. To our knowledge, several recent studies found that ZIKV NS5 facilitates inflammasome to induce IL-1β secretion (66). JEV NS3 interacts with Src protein tyrosine kinase promote inflammatory cytokine secretion (50). DENV NS1 has been reported to interact with STAT3 that enhance the secretion of TNF-α and IL-6 (18). Hence, the non-structural proteins of Flaviviridae play an important role in inflammation response, but the specific signaling pathway and target sites of JEV non-structural protein remains unknown.

Regardless of species of origin, microRNAs (miRNA) are small, noncoding RNAs containing ∼22 nucleotides (nt) that control the posttranscriptional gene regulation by targeting mRNAs. In animal cell, following transcription by RNA polymerase II (RNA Pol II), the RNA polymerase III (RNA Pol III) Drosha in the nucleus process the primary transcripts of miRNAs (pri-miRNAs) into ∼60-100 nt precursors (pre-miRNAs). The pre-miRNAs are then shuttled into cytoplasm, and further processed by RNA Pol II Dicer into ∼22 nt double-stranded (ds) RNA product containing the mature miRNA guide strand and the passenger miRNA strand. The mature miRNA is then load into the RNA-induced silencing complex (RISC) and bind 3’ untranslated regions (UTRs) of target mRNA, which lead to translation repression and/or mRNA degradation. Through the repression of target, miRNA modulate a broad range of gene expression programs during development, immunoreaction and in pathogen infection. Accumulating evidences also suggest abundant of cellular miRNAs involved in multiple stages of the JEV replication cycle. For instance, miR-155, miR-15b miR-19b and miR-29b are induced and activated innated immune response after JEV infection, respectively (3, 60, 61, 73). Beside JEV infection reduces the expression of miR-33a and miR-432 to facilitate virus replication (14, 53). Therefore, the regulation of miRNA is important for cell biological process and mRNA homeostasis during the JEV infection. In contrast to the biosynthetic of miRNA, under physiological or pathological conditions, some of the miRNA decayed rapidly. Most studies on degradation of miRNA have demonstrated the mechanisms involved sequence organization, ribonuclease activity, transcription rates and energy metabolism. For example, several 3’ to 5’ or 5’ to 3’ miRNA-degrading ribonuclease were found to perform degradation of different miRNA in variety of cell types (9, 30). A sequence target-dependent mechanism has also been identified, in which cleavage is mediated by miRNA and high complementary interaction, such as viral non-coding transcript (8, 41), miRNA and miRNA hybrid (13) or extent of sequence complementarity (2). Although the miRNA function has been well defined during the viral replication, the turnover dynamics of miRNA and the mechanisms involved have been poorly defined.

Numerous studies of virus in regulating host miRNA to control cellular protein expression, suggesting a critical role of host miRNAs in immune evasion and viral replication cycle. Both DNA and RNA viruses have developed several mechanisms to degrade or promote cellular miRNA to benefit viral infection and replication. Specifically, herpesvirus saimiri (HVS) and murine cytomegalovirus (MCMV) directly degrade host miR-27 using a virus encode ncRNA which contains miR-27 sequence-specific binding site (8, 39). Similarly, human cytomegalovirus (HCMV) decrease the mature miR-17 and miR-20a using a 15 kb microRNA (miRNA) decay element in the UL/b’ region of the viral genome (37). On the other hand, vaccine virus (VACV) degrades the host miRNA via adding tails using viral poly(A) polymerase (4). Furthermore, the Kaposi’s sarcoma-associated herpesvirus (KSHV) repress the MCPIP1 to facilitate its own miRNA expression (29). Although the direct interaction between host miRNAs and viral components remain far behind other studies, to date, there are two approaches to predict RNA and protein interaction. One of the most cost-effective approaches is computational predicting methods, which could predict protein-RNA interaction and protein-RNA binding sites. Given the limited accurate of computational methods, the RNA-protein immunoprecipitation provides a specific and accurate predication of RNA-protein interaction. Based on those approaches, those would be especially valuable to mapped miRNA-protein binding sites and characterized biological function of viral components in viral replication process.

In the present study, JEV NS3 has been demonstrated to directly disrupt cellular pre-miRNA and reduce mature miRNA levels. Through computational and RNA immunoprecipitation (RIP) analysis, the miRNA binding and turnover of NS3 was found to be arginine-rich domains dependent, which was determined by R226G and R202W of NS3. In addition, NS3 mediated miRNA degradation is critical in inflammation related pathway, which may promote an irreversible inflammatory response leading to neuronal cell death. These results also provide further insight into the role of flavivirus helicase in the regulation of host miRNA turnover.

## Results

### JEV mediates down regulation of global miRNA in mice nervous system

To examine the host miRNAs expression profiles that regulated by JEV infection, miRNA microarrays were used to assay host miRNA expression profiles in mice brain with infection of pathogenic strain JEV (P3). The miRNA expression profiles showed that the abundance of host miRNAs was altered during JEV infection (Fig. 1A). Notably, the 76.7% of total 41 miRNAs (p-value <0.05 and FC > 2.0) were reduced during JEV infection in mice brain. To determine whether the global down regulation of endogenous miRNAs was biased in any way, we used a small RNA deep sequence to analysis the miRNA expression profile of P3 infected NA cells at MOI of 1 and 0.1 for 48 hours (Fig. 1B). This analysis further confirmed that JEV infection led to global host miRNAs decrease. Furthermore, the virus titer and NS3 expression level of cells infected with P3 at 1 MOI was higher than cell infected with P3 at 0.1 MOI, and 1 MOI infection more significantly decreased miRNA expression than 0.1 MOI infection, suggesting that the JEV induced the global miRNA decrease is related to JEV replication (Fig. 1C and D). The miRNA profiles were also confirmed by qRT-PCR analyses in JEV infected NA, BV2 and bEnd.3 cells (Fig. 1E). The integrative analysis of miRNAs and mRNAs was used to predict the major target of the decreasing miRNAs. A total of 42 interacting proteins with 177 interactions were retrieved from the STRING database and 8 proteins were segregated to one subgroup, of which were related to immune and inflammation process (S1 Fig). The result indicated that IL-1β is in the middle of these influential genes and the miR-466d-3p is the only miRNA that decreased significantly (FC ≥ 2.0, p value < 0.01) which target the IL-1β.

**FIG 1.**
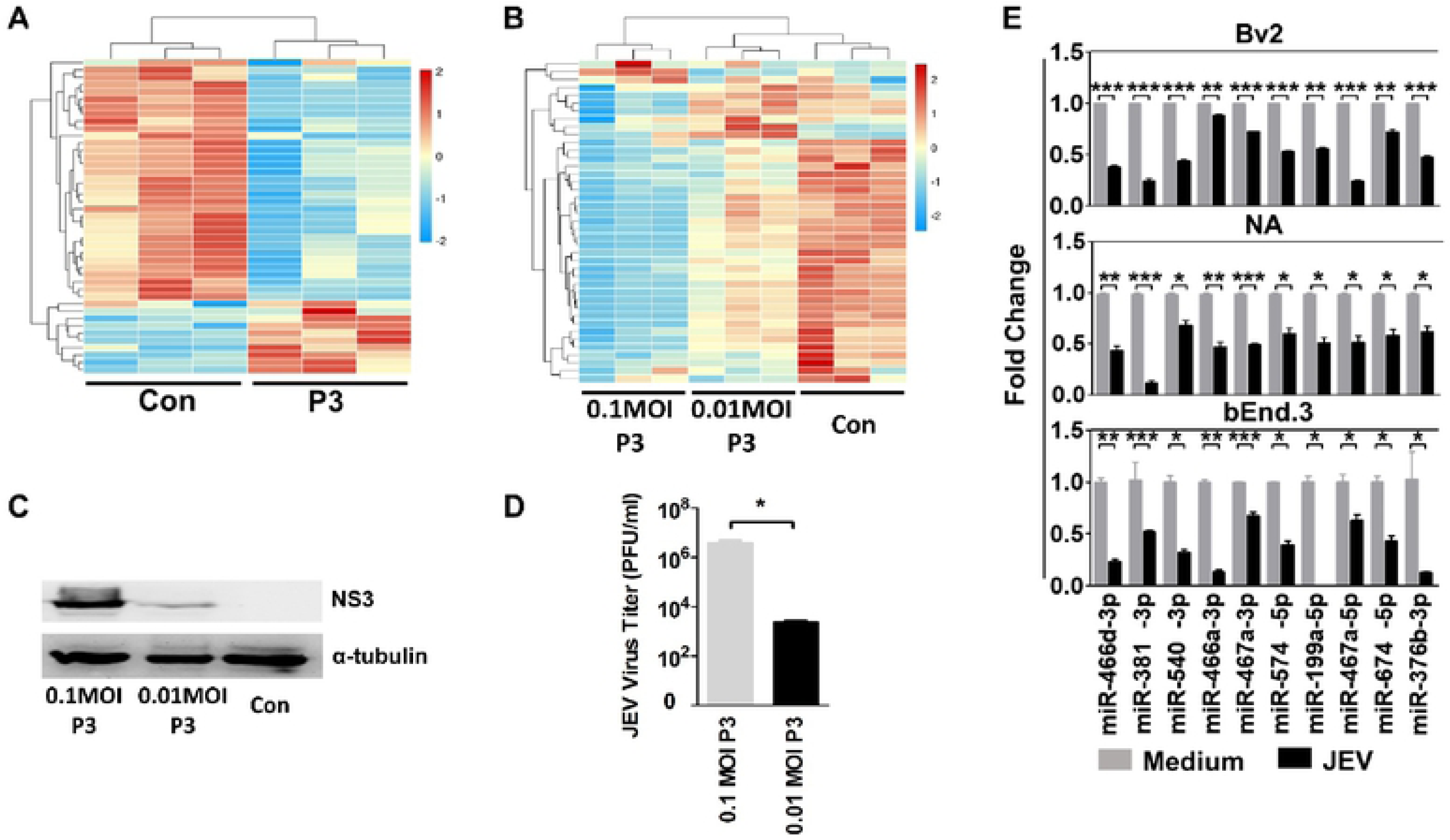
JEV infection downregulates global miRNA expression in mice nervous system. (A) Changes in miRNA expression upon JEV infection of mice brain. The 4-week-old BALB/c mice was infected with 10^3^ PFU of P3 or medium control. After 9 days infection, the miRNA expression level of JEV-infected mice was compared with those from the non-infected control. The color scale is based on log2 changes in expression. (B) Changes in miRNA expression upon JEV infection of NA cells. The NA cells was infected with 0.1 MOI and 0.01 MOI of P3 or medium control. After 48 hours infection, the miRNA expression level of JEV-infected cells was compared with those for the non-infected control. The color scale is based on log2 changes in expression. (C) Western blot of NA cells infected with 0.1 MOI and 0.01 MOI of P3 or medium control after 48 hours. (D) Virus titer of NA cells infected with 0.1 MOI and 0.01 MOI of P3 after 48 hours. (E) Quantitative RT-PCR (qRT-PCR) analyses of the mature miRNA levels in the BV2, NA and bEnd.3 cells after 48 hours JEV infection (1 MOI). For all graphs, results are shown as mean ± SD. Significance was assessed using a Student’s t test, *p≤0.05, **p ≤0.01 and **p ≤0.001.

### JEV infection induces incorrect processing of host miRNAs

Interestingly, the mature miRNA sequence-specific reads were determined by deep sequencing of the 18-24 nt fraction and analyzed by the miRDeep2 module (24), which revealed that percentage of incorrect sequences in the total reads of miR-466d-3p, miR-381-3p and miR-466a-3p were increased (S2 Fig). Furthermore, we observed some of miRNAs from the same precursor were all down-regulated, such as, the miR-466h-3p and miR-466d-3p generated from miR-466d-3p precursor (Fig. 2A), which were both significantly down-regulated in NA cells (Fig. 2B). Notably, the pre-mir-466d-3p and mature miR-466d-3p were reduced in a dose dependent, but the pri-miR-466d-3p appeared unmodified (Fig. 2B). To further examine the miRNA degradation inducing by JEV, the JEV significantly decrease the abundance of miR-466d-3p that derived from an exogenously constructed miRNA mimic compared to the medium control in NA cells (Fig. 2C). Therefore, these experiments indicated that JEV infection decreases the level of pre-miRNA to downregulate miR-466d-3p expression.

**FIG 2.**
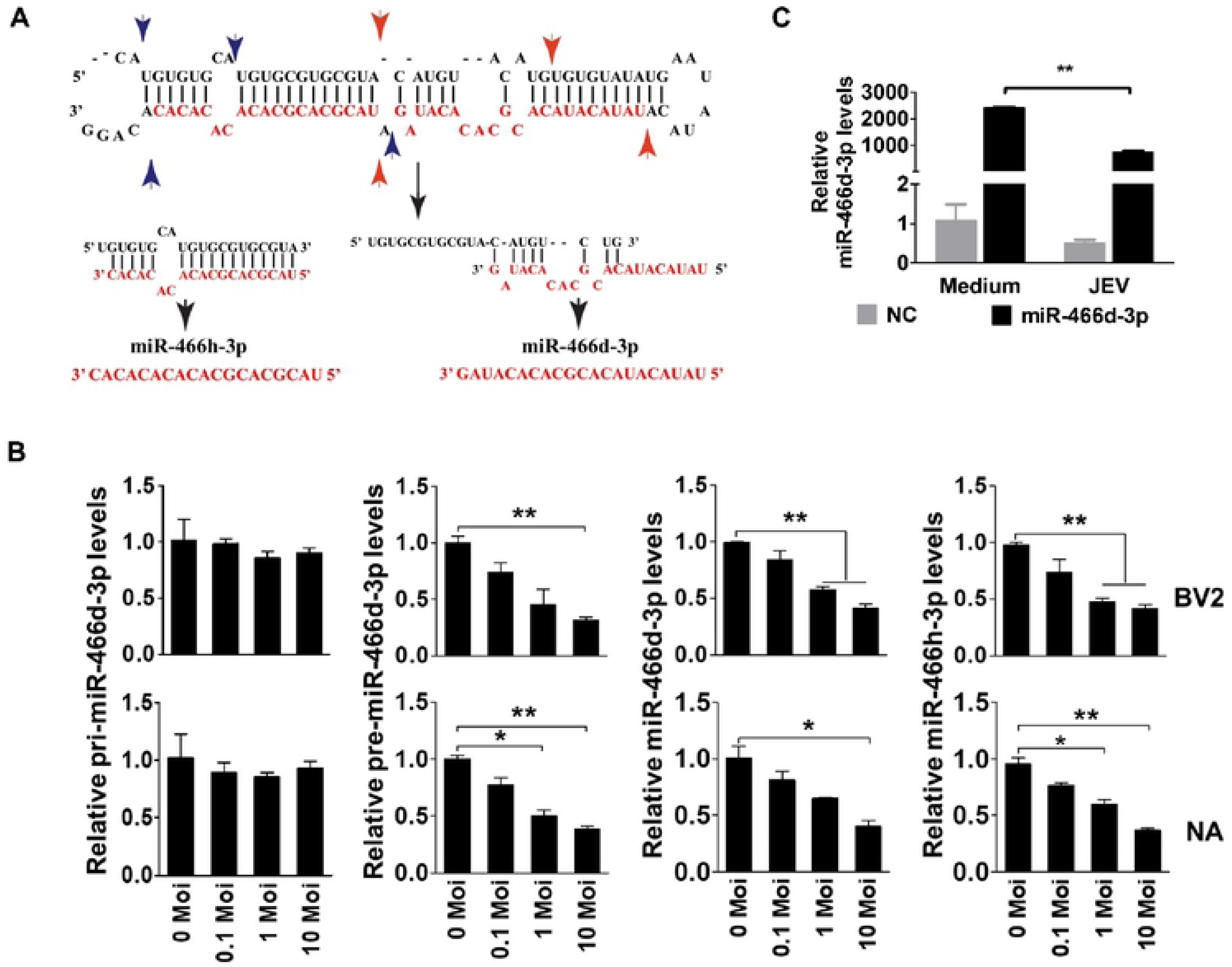
JEV infection induce incorrect processing of host miRNAs. (A) Schematic of pre-miR-466d-3p processing and types of mature miRNAs (miR-466d-3p and miR-466h-3p). The blue arrow represented the miR-466h-3p located in the pre-miR-466d and the red arrow represented the miR-466d-3p located in the pre-miR-466d. (B) Analysis of expression level of mature miR-466d-3p, miR-466h-3p, pre-miR-466d-3p or pri-miR-466d using the quantitative RT-PCR (qRT-PCR) in in NA or BV2 cells at indicated MOI of P3 after 48 hours infection. (C) Quantification of exogenous miR-466d mimic degradation in NA cells by qRT-PCR. The NA cell was infected with P3 at MOI of 0.1 and transfected with miR-466d-3p at 36 hpi. After 48 hours infection, the total RNA from the NA cells was used to quantitative analysis. For all graphs, results are shown as mean ± SD. Significance was assessed using a Student’s t test, *p≤0.05, **p ≤0.01 and **p ≤0.001.

### JEV NS3 mediates degradation of miR-466d-3p in neuronal cells

Expression of the non-structure protein or noncoding sub genomic RNA (sfRNA) of virus is required for viral pathogenicity which would affect the host miRNAs processing (34). JEV infection of neuronal cells results in viral gene transcription and subsequently followed by viral protein translation. The transcriptional inhibitor Favipiravir (T-705) and α-Amanitin or the translation inhibitor cycloheximide (CHX) were used to identify the viral components induced degradation of miR-466d-3p. All early treatments of T-705 and CHX at 12 hours post infection (hpi) could severely block the JEV replication and inhibit degradation of the miR-466d-3p in NA cells. In contrast, late treatment of CHX at 42 hpi could block the degradation of miR-466, but not the treatment of T-705 at 42 hpi (Fig. 3A). Taken together, these data suggested that the posttranscriptional modification or improper processing of host miRNAs is dependent on viral protein production or viral protein with host miRNA interaction in mice neuron cells.

**FIG 3.**
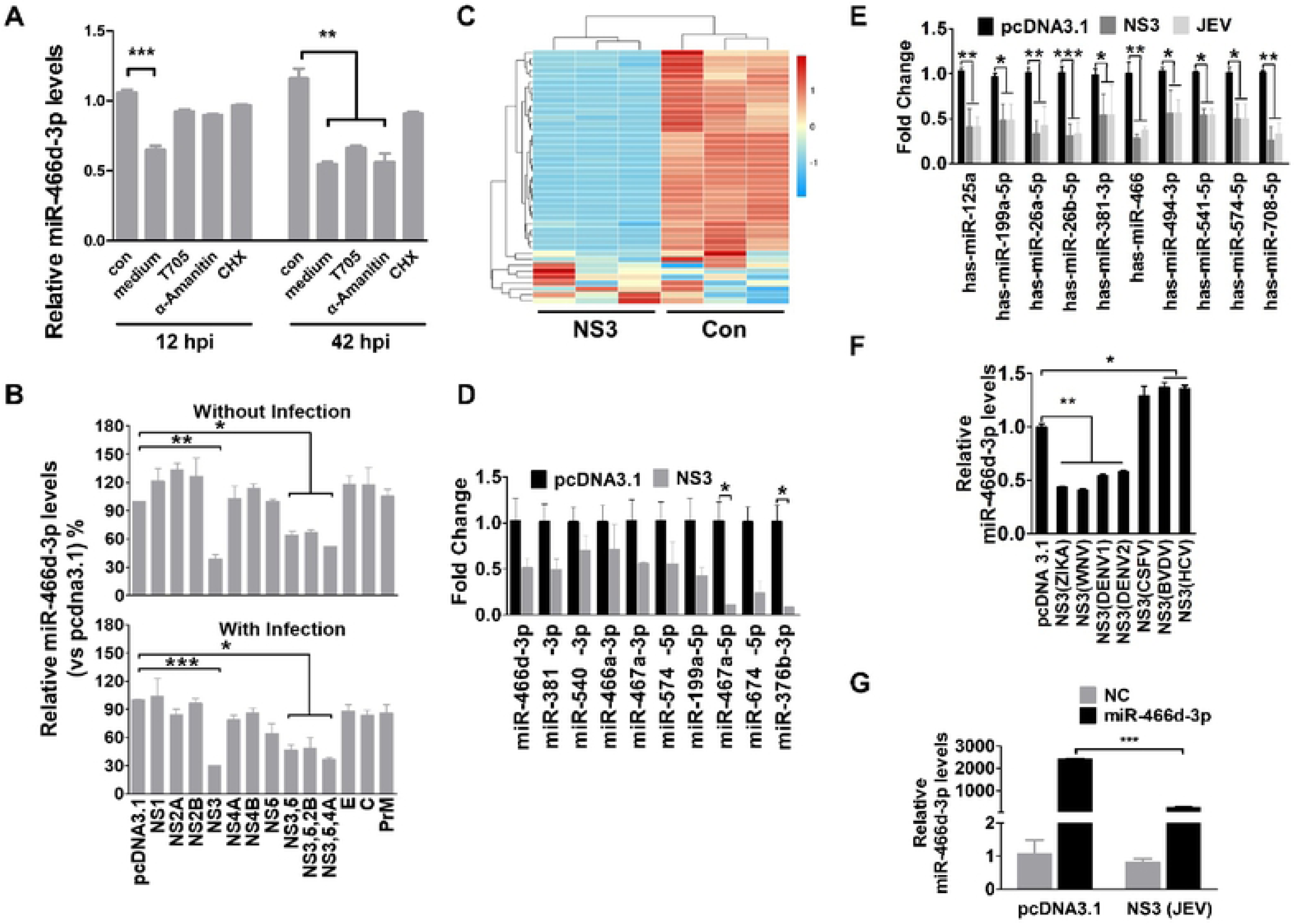
NS3 of JEV-mediated degradation of mir-466d in neuron cells. (A) Transcriptional and translation inhibition assay. The NA cells was infected with P3 at MOI of 0.1 and the T-705, α-Amanitin or CHX were treated with NA cells at 12 hpi or 42 hpi respectively. After 48 hours infection, the total RNA from the NA cells was used to quantify the expression level of miR-466d-3p. (B) Relative analyses of miR-466d-3p expression level in NA cells that transfected with indicated expression plasmid with/without JEV infection. After 48 hours transfection/infection, the total RNA from NA cells was used to quantify the relative expression level of miR-466d-3p (VS pcDNA 3.1 control) by qRT-PCR. (C) Changes in miRNA expression upon transfection of NS3 in the NA cells. The NA cells was transfected with NS3 or pcDNA 3.1 control. After 48 hours transfection, the miRNA expression level of NS3-transfected cells was measured by RNA deep sequencing and compared with those for the pcDNA 3.1-transfected control. The color scale is based on log_2_ changes in expression. (C) qRT-PCR analyses of the mature miRNA levels in NA cells after 48 hours transfection of NS3. (E) The qRT-PCR analyses of the human miRNA in NA cells after 48 hours transfection with NS3 or infection with P3 at 0.1 MOI. (F) The qRT-PCR analyses of the miR-466d-3p in NA cells after 48 hours transfection with NS3 of ZIKA virus, WNV, DENV1, DENV2, CSFV, BVDV and HCV. (G) Quantification of exogenous miR-466d mimic degradation in NA cells by qRT-PCR. The NA cell was transfected with NS3, and after 48 hours the NA cells was transfected with miR-466d-3p mimic. After 54 hours transfection of NS3, the total RNA from the NA cells was used to quantitative analysis. For all graphs, results are shown as mean ± SD. Significance was assessed using a Student’s t test, *p≤0.05, **p ≤0.01 and **p ≤0.001.

Since JEV transcript did not suppress host miRNA, to further determine which viral protein were sufficient to confer the host miRNA degradation, 8 pcDNA3.1 vector encoding each structure protein (E, C and PrM) non-structure protein (NS1, NS2, NS3, NS4A, NS4B and NS5) of JEV were constructed. The miR-466d-3p expression was measured 48 hours post transfection. Only NS3 induced a significant decrease of mature miR-466d-3p (40% reduction compare to pcDNA 3.1 control) (Fig. 3B). Several of other non-structure proteins of JEV have been revolved to associate with NS3 to facilitate the RNA assembling and viral replication (40, 46). To determine the potential association between other different non-structure proteins of JEV, the co-transfection of NS2B, NS4A and NS5 with NS3 were determined and there was no significantly different comparing to NS3 individual transfection group. Moreover, the expression of NS3 in the NA cells also induced a global down regulation of endogenous miRNAs (Fig. 3C) and the same miRNAs in the Fig 1E were also confirmed by qRT-PCR (Fig. 3D). We further assessed the effect of JEV and NS3 in the human neuroblastoma cells (SK-N-SH) to test the regulation of human miRNA by qRT-PCR. As shown in Fig. 3E, both the JEV infection and the NS3 transfection redeuced the miRNA expression level in SK-N-SH cells.

Since the JEV NS3 represented a critical role in turnover of host miRNA, we constructed pcDNA3.1 vector encoding NS3 plasmids of ZIKA virus, WNV, DENV1, DENV2, CSFV, BVDV and HCV to identify whether the helicase of other flavivirus could degrade the host miRNAs. The NS3 of ZIKA virus, WNV, DENV1 and DENV2 induced the decrement of miR-466d-3p when transfected into NA cells (Fig. 3F). However, transfection with NS3 of CSFV, BVDV and HCV were not affected the amount of miR-466d-3p. The alignment of NS3 was revealed that JEV, ZIKA virus, WNV, DENV1 and DENV2 were included in one subgroup and different from the others (CSFV, BVDV and HCV) (S3 Fig). These results indicated that the NS3 of Flavivirus decrease the expression level of miR-466d-3p.

The hypothesis of miRNA degradation and their mRNA-targeting activities upon NS3 expression is not due to inhibiting the expression of RISC. This was partially confirmed by qRT-PCR of the major RISC components gene expression, which of them were all significantly increased after JEV infection (S4 Fig). Furthermore, to explore the possibility that NS3 interacted with dicer or RISC, the proteins that co-immunoprecipitation with NS3 were analysis with LC-MS (S1 Table). The amino sequence identification showed 209 proteins which score were higher than 90 and rich in several functional categories such as, heat shock protein, cytoskeletal components, chaperonin protein and TRiC (TCP-1 Ring Complex). To our knowledge, none of these proteins were correlated with miRNA degradation process that has been reported previously. Furthermore, transfection with NS3 in NA cells significantly decrease the amount of miR-466d that derived from an exogenously constructed miRNA mimic (Fig. 3G), indicating NS3 affect the processing of pre-miRNA or double-stranded segment of miRNA that is 22 bp.

### Block miR-466d enhance IL-1**β** secretion and promote JEV replication

To verify *in silico* analysis of miRNA-mRNA interaction, the level of IL-1β were further determined in JEV-infected mice brain and cells using ELISA. The results revealed that the IL-1β level in mice brain was significantly enhanced at 6 dpi (Fig. 4A). In addition, protein of IL-1β were significantly elevated in the dose-dependent and time-dependent manner in NA, BV2 and bEnd.3 cells.1 (Fig. 4B).

**FIG 4.**
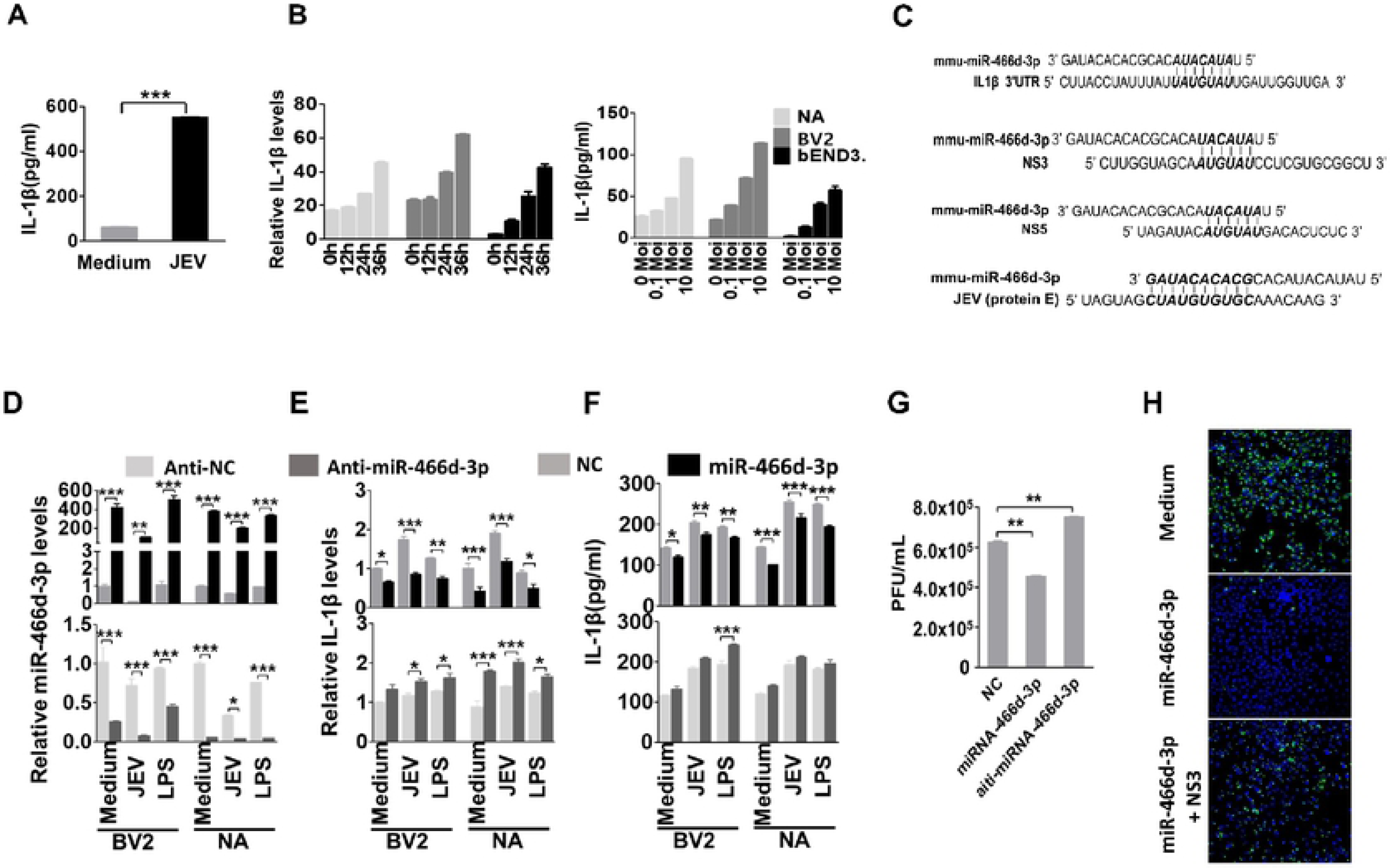
Block miR-466d enhance IL-1β secretion and promote JEV replication. (A) The homogenization of mice brain was collected at 9 dpi and the expression level of IL-1β in mice brain was determined by ELISA. (B) The cell supernatant from NA cells was collected at indicated infection time point and infection dose the expression level of IL-1β was determined by ELISA. (C) Introduction of miR-466d-3p binding sites in IL-1β, NS3, NS5 and E. The miR-466d-3p complementary sequences to the coding sequence of IL-1β, NS3, NS5 and E are indicated as bold and italic. (D-G) The synthetic mimic of miR-466d-3p decreased the expression of IL-1β and the block the replication of JEV, and the inhibitor of miR-466d-3p has the opposite effect. The NA or BV2 cells was infected/treated with JEV at MOI of 0.01, LPS 100ng/ml or medium control, respectively. After 48 hours infection/treatment, the cells was transfected with mimic of miR-466d-3p or inhibitor of miR-466d-3p. (D) After 6 hours transfection, the total RNA from NA cells was used to quantify the relative expression level of miR-466d-3p (VS negative control) by qRT-PCR. (E) After 6 hours transfection, the total RNA from NA cells was used to quantify the relative expression level of IL-1β (VS negative control) by qRT-PCR. (F) After 6 hours transfection, the supernatant of cells was collected to quantify the expression level of IL-1β (VS negative control) by ELISA. (G) After 24 hours transfection, the supernatant of cells was collected to determine the titer of JEV. (H) The miR-466d target sites-fused GFP was co-transfected with NS3 or/and miR-466d-3p mimic, after 48 hours transfection the cells were stained with DAPI. The fluorescence was observed under the fluorescence microscope. For all graphs, results are shown as mean ± SD. Significance was assessed using a Student’s t test, *p≤0.05, **p ≤0.01 and **p ≤0.001.

The sequences of miR-466d-3p and its target site in the 3’ UTR of IL-1β were aligned using TargetScan (http://www.targetscan.org/mmu_72/) (Fig. 4C). To determine whether IL-1β mRNA is indeed target for miR-466d-3p, the expression of IL-1β was examined in BV2 and NA cells after transfected with miR-466d-3p mimics or miR-466d-3p inhibitors, respectively. The results shown that overexpression of miR-466d-3p significantly suppressed IL-1β mRNA and protein levels in BV2 and NA cells, and the similar results were observed in BV2 and NA cells after infected with JEV or treated with LPS (Fig. 4D-F, top). In contrast, miR-466d-3p inhibitor was significantly increased IL-1β mRNA expression and protein secretion in BV2 and NA cells, respectively (Fig. 4D-F, bottom). Moreover, the miR-466d-3p expression level of the cells treated with LPS or poly(I:C) was similar to the medium treated cells and significantly higher than JEV infected cells. However, the miR-466d-3p mimic could still inhibit the LPS or poly(I:C) induced IL-1β over-transcription and overexpression, which indicated that miR-466d-3p is a common regulator of IL-1β not specific to the JEV infection. Interestingly, a miRNA and JEV viral genomic gene comparison analysis indicated that the miR-466d-3p also targets the coding sequence of JEV E, NS3 and NS5 genomes (Fig. 4C). There was a substantial reduction of virus titer in the NA cells that were transfected with miR-466d-3p. Moreover, inhibition of miR-466d-3p leads to an increment of JEV replication (Fig. 4G). To determine whether the miR-466d-3p direct target the seed sequence of JEV, we constructed a miR-466d target sites-fused GFP expressing vector which containing 2 miR-466d target seed sequence of JEV NS3 and E. As expected, transfection of miR-466d mimic resulted in a complete loss fluorescence in miR-466d target sites-fused GFP expression cells, whereas co-transfection of NS3 significant recovered expression of GFP, suggesting that NS3 cleave miR-466d to block the miRNA function (Fig. 4H). Thus, these data suggested IL-1β as a possible target of miR-466d-3p which also worked as a negative regulator of JEV replication.

### Unwinding of miR-466d-3p by NS3 blocks the silencing function of miRNA

The NS3 of JEV previously has been identified as an RNA helicase that acts by unwinding of dsRNA (63). The transactivating response RNA-binding protein (TRBP) and Argonaute 2 contain several dsRNA-binding domains, which facilitate the dsRNA fragments into RISC in order to target mRNA (15, 64). We hypothesized that host pre-miRNAs may also be unwound into a single strand RNA by NS3, in which the RISC could not recognize the single strand fragment or target the mRNA. In vitro unwinding assays reveled that a synthesized pre-miR-466d-3p containing a hairpin structure (Fig. 5A, top) was unwound into a single strand RNA in a dose dependent manner (from lane 2 to lane 5 of Fig 5B). To determine whether NS3 can directly unwind miR-466d-3p mimic in addition to endogenous host miRNAs, using in vitro degradation assay, the synthetic double strand miR-466d-3p mimic (Fig. 5A, bottom) could also unwind into a single strand by NS3 in a dose dependent manner (from lane 2 to lane 5 of Fig. 5C).

**FIG 5.**
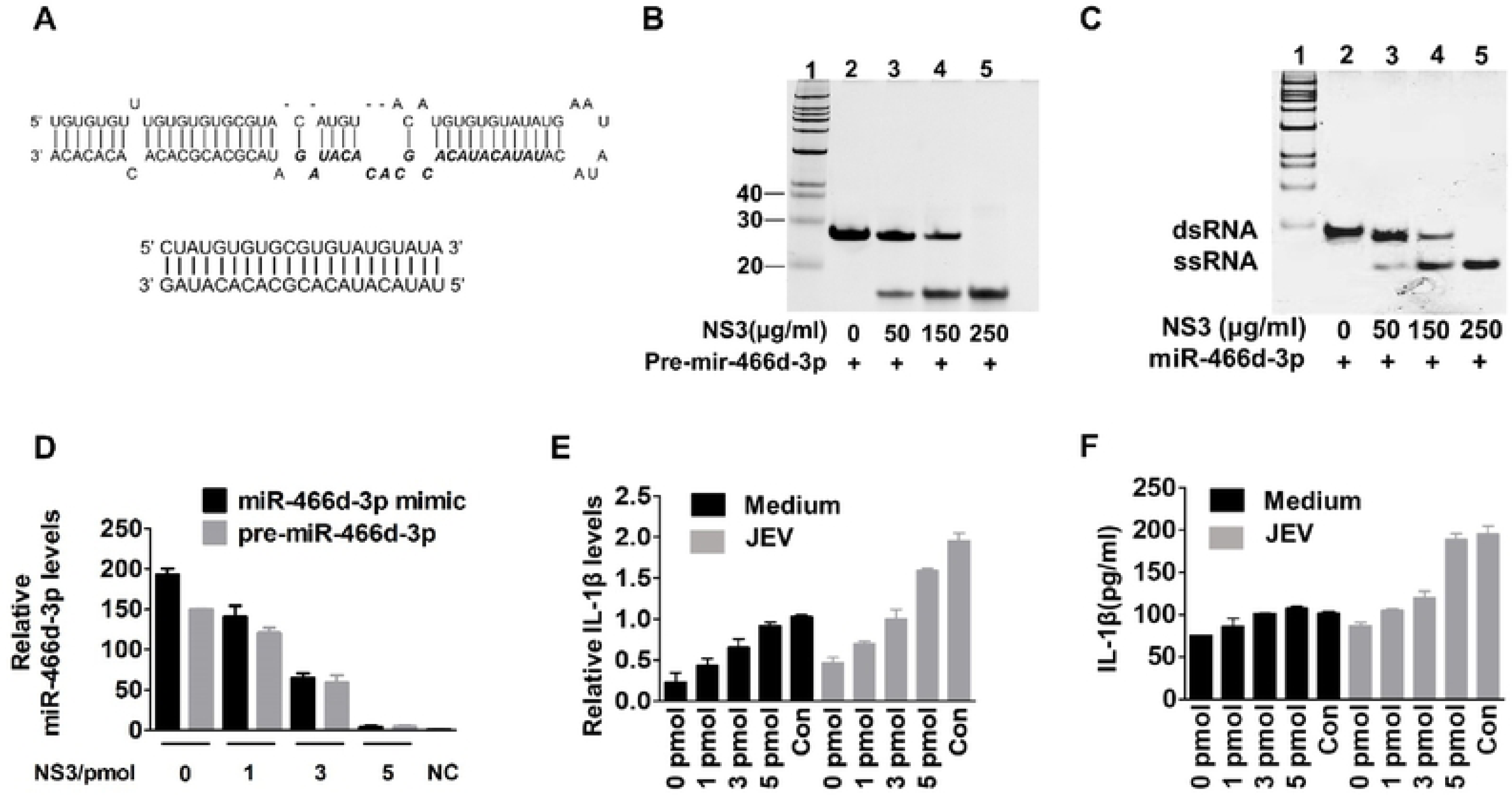
Unwinding of host miRNA by NS3 block the silencing function of miRNA. (A) Schematic of the synthetic double strand pre-miR-466d-3p mimic and double strand miR-466d-3p mimic. The sequence of miR-466d-3p in pre-miR-466d-3p was highlighted in bold. (B-C) The synthetic double strand pre-miR-466d-3 and double strand mimic of miR-466d-3p were unwound by purified NS3 generated from NA cells. The flag-tagged NS3 was expressed in the NA cells and purified by the affinity gel. The indicated concentrations of NS3 (0, 1, 3 and 5 μg/ml) was incubated with 20 pmol of miR-466d-3p mimic (B) or pre-miR-466d-3 mimic (C). After 2 hours incubation at 37℃, the miR-466d-3p mimic or pre-miR-466d-3 mimic was determined by 10% native polyacrylamide gel electrophoresis. (D) The miR-466d-3 mimic and pre-miR-466d-3 mimic were degraded after the unwinding with NS3. After 24 hours transfection, the total RNA of NA cells was used to quantify the relative expression level of miR-466d-3p (VS negative control) by qRT-PCR. (E-F) The unwinding miR-466d-3p mimic by NS3 could not decrease the expression of IL-1β. The total RNA of NA cells was used to quantify the relative expression level of IL-1β (VS negative control) by qRT-PCR (E) and the supernatant of NA cells was used to determine the expression level of IL-1β by ELISA (F). For all graphs, results are shown as mean ± SD. Significance was assessed using a Student’s t test, *p≤0.05, **p ≤0.01 and **p ≤0.001.

To further examine whether the unwinding miRNA could still transport into RISC and execute a normal function of RNA silencing with host mRNA. The fragment of unwinding miR-466d-3p mimic and pre-miR-466d-3p were recycled with Trizol after the in vitro unwinding assay. After transfection with unwinding miR-466d-3p mimic, the expression level of mature miR-466d-3p was similar to the NC control group, indicating that the unwinding miR-466d-3p mimic was unstable in the host cells (Fig. 5D). Notably, we also did not observe the mRNA expression and protein secretion level of IL-1β decreased after transfection of unwinding miR-466d-3p mimic with or without JEV infection (Fig. 5E and 5F). Taken together, these data demonstrate that the NS3 of JEV could facilitate the disfunction of miR-466d-3p in the host cells.

### NS3 has specific binding affinity with pre-miRNA

Many VSRs have been reported to block RNAi by binding siRNA or miRNAs (27, 52). Several ribonucleoproteins performed the post-transcriptional regulatory networks that are mediated by RNA-protein interactions (RPIs), and some computational models were developed to help identify the RPIs and predicated the RNA and proteins binding sites (36, 38, 45, 62). To test whether the miRNAs binding ability is required for NS3 inducing miRNA degradation. In the present study, a RPISeq webserver was used to verify those hypotheses (45). The Random Forest (RF) that calculated by RPISeq of indicated miRNAs between each nonstructural protein of JEV was used to evaluate the RPI interaction. The probability threshold of RF used for positive RPIs was higher than 0.5, and only the value of pre-miRNAs or mature miRNAs that interacted with NS3 were all high than 0.5. The mean RF of NS3 with pre-miRNA and mature miRNA was 0.675 and 0.69 respectively, and significantly higher than all the other proteins of JEV (Fig. 6A).

**FIG 6.**
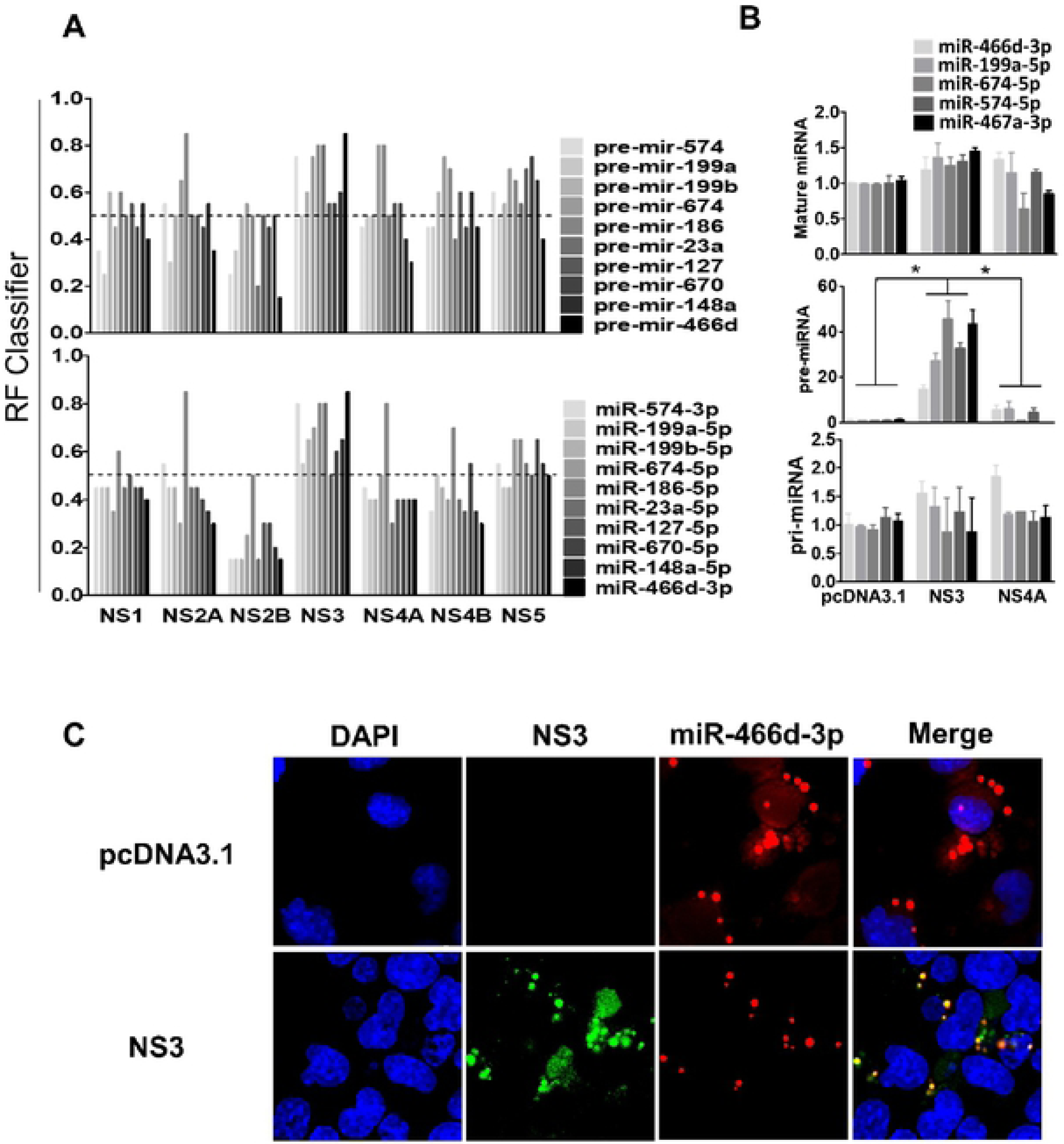
NS3 has specific binding affinity of pre-miRNA. (A) The pre-miRNA and mature miRNA binding affinity of indicated non-structure protein of JEV was predicated with a webserver named RPISeq (http://pridb.gdcb.iastate.edu/RPISeq/) and the Random Forest (RF) that calculated by RPISeq was used to evaluated the RPI. The probability threshold of RF used for positive RPIs was higher than 0.5. (B) The pri-, pre- and mature miRNA level of miR-466D-3p, miR-199a-5p, miR-674-5p, miR-574-5p and miR-467a-3p were selected to determine binding affinity between the NS3 and miRNA by RIP. The NA cells was transfected with plasmid of FLAG-tagged NS3, NS4A or pcDNA 3.1, and these FLAG-tagged proteins was purified by affinity gel after 48 hours transfection. After purification, the total RNA on the FLAG-tagged protein was used to quantify the relative binding affinity of indicated pri-, pre- and mature miRNA (VS pcDNA 3.1 control) by qRT-PCR. The fold change of each RIP reaction from qRT-PCR data was calculated as Fold enrichment=2^(-△ △ Ct[Experimental /pcDNA 3.1])^, △ △ Ct[Experimental/pcDNA 3.1]= △ Ct[Experimental]- △ Ct[pcDNA 3.1], ΔCt=Ct[RIP]-(Ct[Input]-log2(Input/RIP dilution factor)). (C) The pre-miR-466d-3p mimic and NS3 were colocalized in the NA cells. After 48 hours transfection, the CY3 labeled miR-466d-3p mimic and NS3 stained with FITC in NA cells were detected by fluorescence microscope. Images are representative of three independent experiments. For all graphs, results are shown as mean ± SD. Significance was assessed using a Student’s t test, *p≤0.05, **p ≤0.01 and **p ≤0.001.

To further confirm these *in silico* analysis, an RNA immunoprecipitation (RIP) assay using flag-fused protein to identify the NS3 and miRNA binding affinity, and the fold enrichment of non-structure protein over pcDNA 3.1 was used to determine the RIP value (see material and methods for details). The NS3 has a significantly higher pre-miRNA (i.e., miR-466d-3p, miR-199a-5p, miR-674-5p, miR-574-5p and miR-467a-3p) binding affinity than pcDNA 3.1. In contrast to the NS3, the miRNA binding ability of NS4A was similar to the pcDNA 3.1. However, in contrast to these *in silico* analysis, the RIP assay revealed that the binding ability of pre-miR-466d-3p on NS3 was more than nonuple higher than miR-466d-3p and there is no significant binding affinity of NS3 difference between the pcDNA3.1 and the NS4A (Fig. 6B). Furthermore, a CY3 labeled miR-466d-3p mimic revealed that it colocalized with the NS3 expression in the cytoplasm of NA cells (Fig. 6C). However, the miR-466d-3p mimic distributed evenly in the NA cells without the expression of NS3. These results indicate that NS3 specifically binds to pre-miRNA and the binding ability may be essential for miRNA degradation.

### NS3-mediated the miRNA degradation dependent on arginine rich motifs

Computation methods have been reported to predict the amino acid binding sites of PRIs. However, sequence-based predictors are usually high in sensitivity but low in specificity; conversely structure-based predictors tend to have high specificity, but lower sensitivity (38). In order to combine the advantages of two methods, we used several software to predict RPIs binding sites of NS3. The aaRNA web serve that quantified the contribution of both sequence- and structure-based features, the score of RNA and protein binding specificity was represented as binary propensity (from 0 to 1) (38). The RNA binding sites on NS3 of JEV was analyzed using a PDB format of NS3 (PDB entry 2Z83) with aaRNA, 4 arginine (R202, R226, R388 or R464 respectively) on the NS3 has a higher score (> 0.5) than other amino acid sites (S5 Fig). Furthermore, using two web servers called Pprint and PRIdictor (Protein-RNA Interaction predictor) predicted the amino acid binding sites of PRIs (36, 62), the R202, R226 or R464 that located in the helicase region of NS3 has high miRNA binding ability (S6 Fig and S7 Fig). Similarly, previous studies have reported the arginine as a stronger RNA binding amino acids than others (31). Interestingly, almost all 202, 226, 388 or 464 AA sites on the NS3 of flavivirus are argnine but not the hepacivirus or pestivirus. Thus, this sequence alignment confirms the Fig. 3F that only the flavivirus reduce the miRNA expression. When these arginine sites were mutated into other amino acids, the binary propensity or RF value of R202W, R226G and R464Q were all also drop sharply (Fig. 7A and 7B). To further confirm these *in silico* analysis, we constructed 3 substitution mutant vectors of NS3 named as R202W, R226G and R464Q. To investigate the role of miRNA binding affinity in NS3 inducing miRNA turnover, RIP analysis and miRNA expression level were detected in NA cells with these three mutants or parent plasmids of NS3. As shown in Fig. 7C, comparing with the pcDNA 3.1, the mutants of NS3 also specifically bind to pre-miR-466d-3p, and the pre-miR-466d-3p binding ability of R226G and R202W were significantly lower than wild type NS3 (Fig. 7C). Furthermore, the R226G and R202W significantly reduce the degradation of miR-466d-3p, except for R464Q (Fig. 7D). Thus, the 202 and 226 arginine of the NS3 contributes to pre-miRNA binding and promotes miRNA degradation.

**FIG 7.**
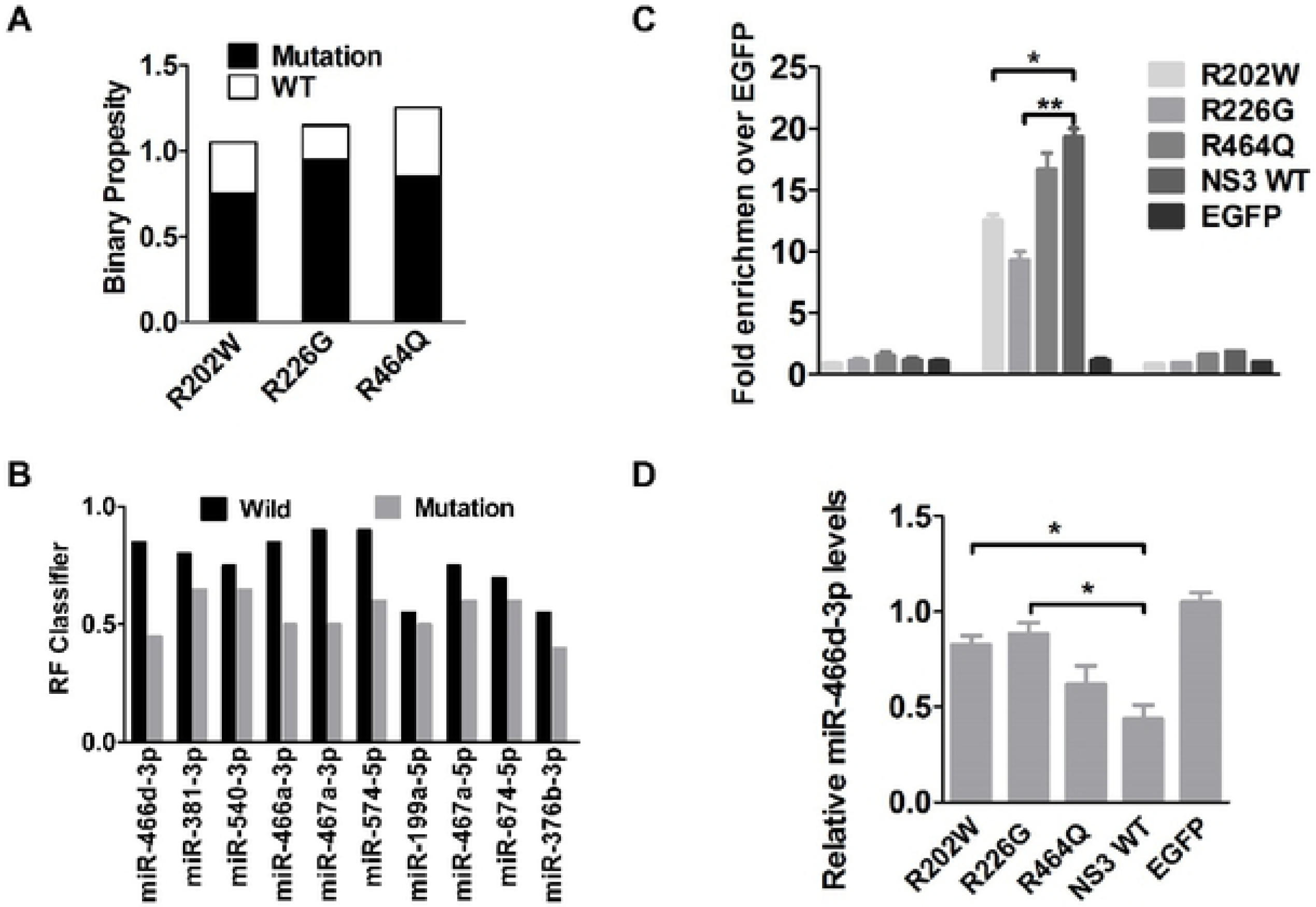
NS3-mediated the miRNA degradation dependent on arginine rich motifs. (A-B) The RNA binding sites on the NS3 were predicted with 3 webservers (supplement) and 3 of arginine has higher RNA binding affinity than others. The sequence of these predicted RNA binding sites was changed to other amino acid as indicated and evaluated the RNA binding affinity by binary propensity (A). After changed all three AA sites (R202W, R226G and R464Q), the probability threshold of RF was evaluated. The probability threshold of RF used for positive RPIs was higher than 0.5 (B). (C) The pri-, pre- and mature miR-466-3p binding affinity of the substitution mutant of NS3 was determined by the RIP assay. Three substitution mutant vectors of NS3 named as R202W, R226G and R464Q was constructed by overlap extension PCR. After 48 hours transfection, the FLAG-tagged NS3 was purified with affinity gel and the total RNA on the NS3 was used to quantify the relative binding affinity of pre-miR-466-3p (VS pcDNA 3.1 control) by qRT-PCR. The fold change of each RIP reaction from qRT-PCR data was calculated as Fold enrichment=2^(-△ △ Ct[Experimental /pcDNA 3.1])^. (D) Relative analyses of miR-466d-3p expression level in NA cells that transfected with indicated expression plasmid. After 48 hours transfection, the total RNA from NA cells was used to quantify the relative expression level of miR-466d-3p (VS pcDNA 3.1 control) by qRT-PCR. For all graphs, results are shown as mean ± SD. Significance was assessed using a Student’s t test, *p≤0.05, **p ≤0.01 and **p ≤0.001.

## Discussion

Increasing studies have shown that miRNAs play an important role in the replication and propagation of viruses, including defense of pathogenic viral infections or promotion of viral replication through complex regulatory pathways (26). In the early steps of viral infection, innate viral detected sensors of host cells, such as pathogen recognition receptors (PRRs), recognize a large spectrum of pathogens and initiated a downstream antiviral pathway activates that include miRNAs (65). However, the defense function of host miRNAs can be counteracted by viral suppressors of RNA silencing (VSRs) that inhibit host antiviral responses by interacting with the critical components of cellular RNA silencing machinery or direct participate the host RNA degradation (6, 26). Recently, an increasing number of studies have found that JEV infection were capable of regulating functional miRNAs (3, 14, 53, 60, 61, 73). Although plenty of works were examined the host biogenesis regulating by miRNA during the viral infection process, relatively little was known about regulation of miRNA by JEV. Given the importance of miRNA in establishing infection of JEV, our research was aim to explore the interplay between the JEV and host miRNAs.

An intriguing aspect of this study is that the JEV globally decrease the host miRNA and independent on Dicer or RISC. We demonstrated that the JEV NS3 could unwind the pre-miR-466d and induce the disfunction of miR-466d-3p. We also observed that the decrement of miR-466d-3p could enhance the JEV replication. Together, these results suggested a role of the miRNA degradation in enhancing JEV replication. Similarly, poly(A) polymerase of vaccina virus (VACV) and 3’UTR of the murine cytomegalovirus degrades the host miRNA via different mechanisms (4, 39). Although this is the first report of a link between the host miRNA degradation and JEV infection, several studies have reported the role of miRNA machineries in the replication of flavivirus, especially dengue virus and Kunjin virus (34, 43). Some analogous systems in degradomes of bacterial and exosomes of eukaryotes are associated with RNA helicases to perform the RNA degradation (22, 49). Hence, the helicase of the JEV seem to be involved in degrading the host miRNA.

Why would JEV has evolved a miRNA degradation processing during viral infection in the CNS? There are two possibility why the JEV infection induced global degradation of host miRNA might promote viral infection and replication. First, the host cell encoding miRNA binds to miRNA-binding sites of viral genomes to influence viral function (56, 72). Consistent with this is our observation that restoring miR-466d-3p expression during the JEV infection reduces viral replication. Conversely, the viral induced degradation of host miRNA leads to block the cleavage of viral genome and viral protein. Second, changes in host miRNA expression could also lead to enhance antiviral effector resulting in reducing viral replication (54, 61). However, some viral infections mediate degradation of host miRNAs leading to downstream changes in host transcriptome that can be benefit virus replication and pathogenicity (4). These hypotheses were also confirmed with NS3 of other Flavivirus, including ZIKA could decrease the expression level of miR-466d-3p to enhance the inflammation.

Eubacteria, Archaea, and eukaryotes have developed dedicated pathways and complexes to check the accuracy of RNAs and feed undesired RNA into exoribonuclease that degrade RNA. Furthermore, these complexes usually contain adaptor proteins such as RraA (a regulator of RNase E and DEAD-box helicases), RhlB (a DEAD-box RNA helicase) (49), Ski2-like RNA helicases (22, 32) and Suv3 RNA helicase (20). In addition to the RNA degradation, the UPF1 helicase can dissociate miRNAs from their mRNA targets and promote the miRNAs degradation by Tudor-staphylococcal/micrococcal-like nuclease (TSN) (21). The P68 helicase promote unwinding of the human let-7 microRNA precursor duplex, which help let-7 microRNA load into the silencing complex. p72 (DDX17) interact with the Drosha-containing Microprocessor complex and facilitate the processing of a subset of pri-miRNAs in the nucleus and miRNA-guided mRNA cleavage (51). Although host helicase may initially seem to benefit only the cells processing and immune system, multiple studies have shown that viral helicase promote viral replication and disrupt immune system. The NS3 of flavivirus are capable of unwinding both the DNA and the RNA that are not of the viral own origin (16, 63). Similar, we observed NS3 of JEV could reduce exogenously constructed ds-miRNA mimic and disable the miRNA mimic.

Furthermore, it has been shown that some single-amino-acid mutation in R538, R225 and R268 of DENV, which affect the replication of DENV (17, 58). Another study report that the Asp-285-to-Ala substitution of the JEV NS3 protein abolished the ATPase and RNA helicase activities (63). Furthermore, the helicase domains of NS3 alone is known to induce cell apoptosis in neuron cells (68, 71). These findings suggested that NS3, particularly helicase domain, contributes to the adaptation of flavivirus for efficient replication. Thus, to investigate the details of the NS3 involvement in miRNA degradation, using miRNA deep sequence, we analysis the subtype sequence of mir-466d during the JEV infection and NS3 over expression, in which the percentage of incorrect splicing products of mature miRNA was higher than normal. Furthermore, the in vitro unwinding assay demonstrated that the NS3 could unwinding the pre-miR-466d and induce the disfunction of miRNA. However, the molecular mechanism of miRNA degradation after unwinding by NS3 need to be clarified in next step experiment.

In the present study, we proved that the arginine of NS3 are critical for host pre-miRNA binding and promote the global host miRNA turnover. We constructed 3 arginine variants of NS3 that have single amino acid substitutions on R202W, R226G and R464Q, which were reduced pre-miRNA binding affinity of NS3 and the degradation of miRNA was almost abrogated. Interestingly, these sites are all located in the helicase region of NS3, which from protein sequence of NS3 163 to 619. Similar to our findings, a number of proteins containing arginine-rich motifs (ARMs) are known to bind RNA and are involved in regulating RNA processing in viruses and cells. Such as the ARM of lambdoid bacteriophage N protein or HIV-1 Rev protein bind RNAs and regulate the RNA transport and splicing (23, 57). In addition, four single amino substitution of arginine on HIV-1 Rev protein strongly decrease their RNA binding ability. In brief, these results extend the helicase of Flavivirus function beyond unwinding duplex RNA to the decay of pre-miRNAs, which was provided a new mechanism of helicases in regulating miRNA pathways. Our results suggested that helicase of flavivirus may have the capacity to regulate various cellular miRNAs, which would be a part of a general viral response to overcome host defense mechanisms.

## Materials and Methods

### Cell and viruses

The mouse microglia cell line BV-2, the mouse neuroblastoma cell line NA and mouse brain endothelial cell line bEnd.3 were gift from Huazhong Agricultural University which maintained in Roswell Park Memorial Institute (RPMI) 1640 medium (Thermo-Fisher, USA) supplemented with 10% fetal bovine serum (FBS; Gibco, Carlsbad, CA, USA) at 37°C in 5% CO2. The baby hamster Syrian kidney cells line BHK-21 was gift from Huazhong Agricultural University which cultured in Dulbecco’s modified Eagle’s medium (DMEM, High glucose, Thermo-Fisher, USA) containing 10% FBS. The P3 strain of JEV (Japanese encephalitis virus) was propagated in suckling BALB/c mice (purchased from Vital River Laboratories, China) as previously (37). Briefly, one-day-old suckling mice were inoculated with 10 μl of viral inoculum by the intracerebral (i.c.) route. When moribund, mice were euthanized and brains removed. A 10% (w/v) suspension was prepared by homogenizing the brain in DMEM and centrifuged at 10,000 g for 5 min to remove cellular debris. The brain suspension was filtered through 0.22 µm-pore-size sterile filters (Millipore, USA) and sub packaged at −80°C until further use.

### RNA sequencing

Total RNA was extracted from culture NA using the Total RNA Extractor Kit (B511311, Sangon) according to the protocols. A total of 2μg RNA from each sample was used for library preparation according to the manufacturer’s instructions of the VAHTSTM mRNA-seq V2 Library Prep Kit for Illumina®. The library fragments were purified with AMPure XP system (Beckman Coulter, Beverly, USA). PCR products were purified (AMPure XP system) and library quality was assessed on the Agilent Bioanalyzer 2100 system. The libraries were then quantified and pooled. Paired-end sequencing of the library was performed on the HiSeq XTen sequencers (Illumina, San Diego, CA). FastQC (version 0.11.2) was used for evaluating the quality of sequenced data. Raw reads were filtered by Trimmomatic (version 0.36) according to protocol. And the remaining clean data was used for further analysis. Gene expression values of the transcripts were computed by StringTie (version 1.3.3b). Principal Component Analysis (PCA) and Principal co-ordinates analysis (PCoA) were performed to reflect the distance and difference between samples. The TPM (Transcripts Per Million), eliminates the influence of gene lengths and sequencing discrepancies to enable direct comparison of gene expression between samples. DESeq2 (version 1.12.4) was used to determine differentially expressed genes (DEGs) between two samples.

### Plaque assay

JEV was titrated on BHK-21 cells line by viral plaque formation assay as previously (14). A monolayer of BHK cells was co-cultured with JEV (a serial 10-fold dilution prepared in DMEM without FBS) at 37°C. After 2 h incubation, the DMEM containing 3% FBS and 4% sodium carboxymethyl cellulose (CMC) (Sigma) were added to the cells and cultured for 5 days. Until the appearance of visible plaques, the cells were fixed with 10% formaldehyde overnight, followed by staining with crystal violet for 2 h. Visible plaques were counted and the viral titers were calculated. All data were expressed as means of triplicate samples.

### Transfection of cells with miRNA mimics or inhibitors

Mouse miR-466d-3p mimics, inhibitors, mimics control and negative controls were purchased from GenePharma (China). The sequences of the mimics, inhibitors, mimics control and controls oligo nucleotides were as follows: miR-466d-3p mimics, 5’-UAUACAUACACGCACACAUAG-3’ (forward) and 5’-AUGUGUGCGUGUAUGUAUAUU-3’ (reverse); mimic negative controls, 5’-UUCUCCGAACGUGUCACGUTT-3’ (forward) and 5’-ACGUGACACGUUCGGAGAATT-3’ (reverse); miR-466d-3p inhibitor, 3’-CUAUGUGUGCGUGUAUGUAUA-5’; and inhibitor negative control, 3’-CAGUACUUUUGUGUAGUACAA-5’. All of miRNA mimics (1.5 µmol/well), inhibitors (1.5 µmol/well) and control (3 µmol/well) were transfected into BV-2 cells or NA cells (10^6^ cells/well) in 12-well plates using Lipofectamine 3000 (Invitrogen) according to the manufacturer’s instructions.

### Viral infection

BV-2 cells, NA cells or bEnd.3 cells were seeded in 12-well plates (10^6^ cells/well). Until grown to 80% confluence, the cells were incubated with serum-free medium or JEV at a multiplicity of infection (MOI) of 0.1 at 37°C for 2h. After discard of unbound virus and medium, the cells were cultured in 1640 with 10% FBS, 1% Penicillin-Streptomycin Liquid (PS) at 37°C in 5% CO2 for 48h. Cells and supernatants were collected 48 h post-infection (hpi).

### RNA isolation and Reverse transcription real-time PCR

The total RNA of the cells was isolated by trizol reagent (Thermo Fisher) according to the Manufacturer’s recommendations and used for qRT-PCR in an Applied Biosystems® 7500 Real-Time PCR Systems (Thermo fisher) as described previously. For the evaluation of mRNA, to obtain cDNA, the total RNA (1 µg) was reverse transcribed by PrimeScript RT reagent Kit with gDNA Eraser (Takara, China). The IL-1β and components of RISC mRNA expression levels of the NA cells were quantified with the SYBR Green qPCR kit (Takara) by following the manufacturer’s instruction using gene-specific primers (S2 Table). Amplification was performed at 95°C for 30 s, 95°C for 5 s, 60°C for 34 s, followed by 40 cycles of 95°C for 15 s, 60°C for 1 min, 95°C for 30 s, and 60°C for 15 s. The expression of IL-1β was normalized to the level of glyceraldehyde-3-phosphate dehydrogenase (GAPDH) expression using the 2-△△CT method as previously (69). For the expression of miRNA, the reverse transcription primers of all miRNA were added with specific stem-loop structure at the end of 5’ (S3 Table). The pri-, pre- and mature-miRNA levels of cells were quantified with the SYBR Green qPCR kit using miRNA sequence-specific primers (S4 Table and S5 Table). Amplification was performed at 95°C for 30 s, 95°C for 5 s, 60°C for 34 s, followed by 40 cycles of 95°C for 15 s, 60°C for 1 min, 95°C for 30 s, and 60°C for 15 s. The relative levels of microRNAs were normalizing to U6 determined by the 2-△△CT method as previously (73).

### ELISA

The culture supernatants of the experimental group and control group cells or brain tissue lysates were collected and stored at −80°C. The protein levels of IL-1β in cell cultures or mouse brain tissue lysates were determined by enzyme-linked immunosorbent assay (ELISA) kits (eBioscience) according to the manufacturer’s instructions.

### Immunoprecipitations

Immunoprecipitation with FLAG fused protein was performed as described previously (70). Briefly, BSR cells was transfected with the FLAG-tagged plasmid. At 48 hpi, the cells expressing FLAG fusion protein were harvested and lysed with CelLyticTM M lysis buffer (Sigma) containing protease inhibitor cocktail (Sigma) in 4°C for 30 minutes on a shaker at 10 rpm. Each cell lysate was incubated with ANTI-FLAG M2 affinity gel (Sigma) at 4°C overnight on a shaker at 10 rpm. Then, the agarose gel was centrifuged for 30 seconds at 8200 g to remove the supernatants. After three times washed with 0.5 ml of TBS, the bound proteins were eluted by boiling with SDS-PAGE loading buffer for 5 min and determined with Western blotting.

### Western blotting

The cells were lysed with RIPA Lysis and Extraction Buffer (Thermo Scientific), and the protein level was measured with the Enhanced BCA Protein Assay kit (Sigma-Aldrich). The extracts which contained 25 μg of total proteins were subjected to 10% SDS polyacrylamide gel, and protein blots were transferred to Nitrocellulose membrane (NC) after electrophoresis. The membrane was then washed with TBST and blocked in 5% skimmed milk at 4°C overnight. All primary antibodies were prepared at a dilution of 1:1,000 in 1% bovine serum albumin (BSA) in 1×PBST, followed by horseradish peroxidase-conjugated secondary antibodies (Sigma) for 1h at room temperature. Blots were detected by enhanced chemiluminescence reagent (Thermo Scientific) and developed by exposure in Tanon 5200 (Tanon) using Tanon MP software. In addition, α-tubulin played the role of an internal control.

### RNA immunoprecipitation (RIP)

RIP was performed as previously (48), with slight modification. Briefly, NA cells was transfected with the FLAG-tagged NS3, FLAG-tagged NS2B or FLAG-tagged pcDNA 3.1 pcDNA3.1 plasmid for 48h. 10^7^ cells expressing FLAG fusion protein were harvested with 2.5% trypsin and resuspended in 5ml PBS. 143μl of 37% formaldehyde was added to the resuspension to cross-link for 10 minutes on a shaker, and 685μl of 2Mthe glycine was used to block the formaldehyde. The cells were washed twice by 5ml ice-cold PBS and centrifuged for 2 minutes at 400 g to collect the cells pellet. 1ml CelLytic^TM^ M lysis buffer (Sigma) containing 20μl 0.1M Phenylmethylsulfonyl fluoride, 20μl protease inhibitor cocktail (Sigma) and 5μl 40U/μl RNase inhibitor (Invitrogen) was added to cells, and the cells were kept on ice and sonicated for 2 mins (10 sec on, 10 sec off, Amplitude 15 μm) until the lysate is clear.

Followed by centrifuging the lysate for 3 minutes at 14000g to collect supernatant. Each 1ml cell lysate was added to the 40μl washed with ANTI-FLAG M2 affinity gel (Sigma) at 4°C for 4h on a shaker at 10 rpm. The next step was to centrifuge the resin for 30 seconds at 8200 g and remove the supernatants. After the resin was washed three times with 0.5 ml 500μl of TBS, the total RNA was extracted with Trizol reagent and analyzed by RT-qPCR.

The fold change of each RIP reaction from RT-qPCR data was calculated via 2-△△CT method as previously with minor modification (42) and the formulas see below. All the Ct value of each specimen (FLAG-tagged NS3, FLAG-tagged NS2B or FLAG non-fused blank pcDNA3.1) was normalized with the Input RNA to eliminate the possible differences in RNA samples preparation (△Ct). To obtain the △△Ct, the normalized experimental RIP fraction value (△Ct) was normalized to unspecific background as an internal control (ΔCt normalized of pcDNA3.1sample). Finally, the △△Ct [Experimental/pcDNA3.1] was performed with linear conversion to calculate the Fold enrichment. The formula of Fold enrichment is shown below.

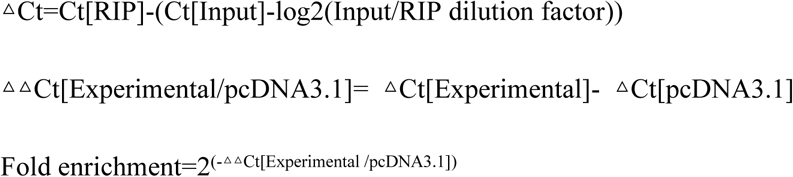

### MicroRNA target prediction

The sequences for the miRNA whose expression changed during JEV Infection have been submitted to the public miRNA database miRBase (www.mirbase.org). The miR-466d-3p target binding sites in the 3’UTRs of mouse gene transcripts were identified with Target Scan software (http://www.targetscan.org/).

### Bioinformatics Analysis of the changed miRNA

The total RNA was isolated from mouse brains with Trizol reagent (Invitrogen) for miRNA and mRNA Microarray. miRNA and mRNA hybridization were performed by shanghaiBio Corporation (shanghai, China) with the use of 8 × 15 K Agilent Mouse microRNA Microarray and 4 × 44 K Agilent Whole Mouse Genome Oligo Microarray. For each sample pair, the experiments were done with two independent hybridizations for miRNA (Agilent’s miRNA Complete Labeling and Hyb Kit) or for mRNA (Cy3 and Cy5 interchanging labeling). Hybridized arrays were scanned at 5 μm resolution on a Microarray Scanner (Agilent p/n G2565BA). Data extraction from images was done by using Agilent Feature Extraction (FE) software version 9.5.3. The mRNAs which were caused significant changes (change fold ≥ 2.0, p value < 0.05) by JEV infection in mouse brain were clustered using GO and KEGG enrichment tools (ShanghaiBio Analysis System). And the interaction of the most significantly differential expression proteins was retrieved by STRING (59).

### Construction of mutant NS3 plasmids

The plasmid encoding FLAG-tag E, C, PrM, NS1, NS2A, NS2B, NS3, NS4A, NS4B, and NS5 cDNA clone in pcDNA3.1 (+) flanked by Kpn I ribozyme and Xba I ribozyme sequences as described previously (70). The AA residues at R202W, R226G R464Q of NS3 in the SA strain were swapped individually or both by overlap extension PCR as described previously. Briefly, in a PCR reaction 1 μl of full-length NS3 gene cDNA was mutated and amplified with 20 μM each of 202, 226 or 464 site mutation forward and reverse primers and 20 μM each of JEV-NS3 forward and reverse primers using PrimeSTAR GXL DNA Polymerase (TAKARA) according to the manufacturer’s instructions. The size of PCR products was 1.86 kb and the products were purified by gel purification kit. The PCR mixture was heated at 94 °C for 2 min, followed by 35 cycles of amplification at 98 °C for 10 s, 55 °C for 30s and 68 °C for 1min45s, and a final extension at 68 °C for 10 min. All NS3 fragments with FLAG-tag that included AA mutations and NS3 cDNA vector were digested with enzyme sets KpnI and XbaI (Thermo Scientific). Following the digestions, the NS3 fragments with FLAG-tag and NS3 cDNA vector were ligated together at an approximate molar ratio of 1:3 using TaKaRa DNA Ligation Kit LONG (TAKARA) according to the manufacturer’s instructions.

### The integrative analysis of miRNAs and mRNAs

Usually, there were more than hundreds target genes among each miRNA used 3 miRNA target predication databases TargetScan (1), miRDB (67) and microRNA (5). The integrative analysis of miRNAs and mRNAs allowed us to predict the major target of the decreasing miRNAs. To accurately elucidate the correlation between the mRNA expression pattern and miRNA targeted regulation, mRNA expression files were measured with mRNA microarray to predict the major decreasing miRNAs in mice brain. Firstly, those abnormal miRNAs target mRNAs were used to find out the common mRNA (with a threshold of fold change ≥ 2.0, p value < 0.01) from the mRNA expression profile and miRNA expression profile. Using those common genes as seed, a protein-protein interaction network was constructed to discover the core gene of miRNA target. A total of 42 interacting proteins with 177 interactions were retrieved from the STRING database (protein-protein interaction enrichment p value < 1.0e-15). The k-Means clustering algorithm was applied to segregate the network of those interacting proteins into different subgroup.

### LC-MS analysis

The FLAG fused NS3 was purified by immunoprecipitation and washed with pure water twice in 0.5 ml Eppendorf tube. The following procedure to digest was performed as previously described (33). To dehydrate, the specimen was treated with 50% acetonitrile for 30 min send following with 100% acetonitrile for another 30 mi. After dehydration, the gel was dried in SpeedVac concentrator (Thermo Savant, Holbrook, NY, USA) for 30 min then restored with reduction buffer (25mM NH4HCO3 in 10mM DTT) at 57°C for 1 hafted removing reduction buffer, the gel was dehydrated again with 50% and 100% acetonitrile respectively for 30 min. To rehydrate, the gel was removed acetonitrile and incubated with 10μL 0.02μ g/μ L trypsin in 25mM NH4HCO3 for 30 min at RT. For tryptic digestion,20 µL cover solution (25mM NH4HCO3 in 10% acetonitrile) was added for digested 16 hours at 37°C, and the supernatants were transferred into another tube. For peptide extraction, the gel was extracted with 50 µL extraction buffer (5% TFA in 67% acetonitrile) at 37°C for 30min. Finally, the peptide extracts and the supernatant of the gel were combined and then completely dried in Speed Vac concentrator. The specimen was analyzed by the direct nanoflow liquid chromatography tandem mass spectrometry (LC-MS/MS) system and the ion spectra data were identified in the protein database.

### Ethics Statement

The experimental infectious studies were performed in strict accordance with the Guide for the Care and Use of Laboratory Animals Monitoring Committee of Hubei Province, China, and the protocol was approved by the Scientific Ethics Committee of Huazhong Agricultural University (protocol No. Hzaumo-2015-018). All efforts were made to minimize the suffering of the animals.

### Statistical analysis

All experiments were performed at least three times with similar results. The data generated were analyzed using GraphPad Prism 5 (GraphPad Software, San Diego, CA). For all tests, the P value of <0.05 was considered significant.

## Supporting Information Legends

S1 Fig. **The protein interaction network was constructed by STRING that based on the candidate genes (42 genes) of common mRNA from the miRNAs and mRNA expression profile**. Light blue lines, indicated interactions from curated databases. Purple lines, experimentally validated interactions. Green lines, predicted interactions from gene neighborhoods. Red lines, predicted interactions from gene fusions. Dark blue lines, predicted interactions from gene co-occurrence. Yellow lines, represented interactions from textmining. Light purple lines, interactions from protein homology. Blank lines, interactions from co-expression. And the dotted lines indicated interactions of bridge different subnetworks. The colored nodes in the network showed the query proteins and first shell of interactors. The network is divided into many groups according to biological Process (GO). The red nodes in the network represented the genes which played a role in the immune and inflammation process. Average number of connections per node is 2.81 (protein-protein interaction enrichment p value < 1.0e-15).

S2 Fig. **The mature miRNA sequence-specific reads were determined by deep sequencing of the 18–24 nt fraction of JEV-infected or NS3-expressed cells.** The mature miR-466d-3p, miR-381-3p, miR-466a-3p, miR-467a-3p, miR-199a-5p and miR-466h-3p sequence-specific reads were determined by deep sequencing of the 18-24 nt fraction of P3-infected cells, or NS3 transfected cells. The bold type sequence represents the correct sequence of the miRNA. The percentage of indicated sequence reads was analyzed by the miRDeep2.

S3 Fig. **Phylogenetic tree and alignment based on the full-length NS3 sequences of 37 Flaviviride virus strains.** Phylogenetic tree deduced from a full-length alignment of NS3 from indicated viruses using the neighbor-Joining method as implemented in MEGA7 (Left). The numbers below the branches are bootstrap values for 1000 replicates. Phylogenetic analysis of 37 NS3 gene nucleotide sequence from Flaviviride included JEV vaccine strain SA-14-14-2 (GenBank No. M55506.1), lineage II WNV strain (GenBank No. M12294.2), Dengue virus 1 isolate TM100 (GenBank No. KU666942.1), Dengue virus 2 isolate TM26 (GenBank No. KU666944.1), Zika virus strain MR 766 (GenBank No. AY632535.2), Classical swine fever virus C-strain (GenBank No. Z46258.1), Chikungunya virus isolate MY/08/4567 (GenBank No. FR687343.1) Hepatitis C virus QC69 subtype 7a (GenBank No. EF108306.2) and Bovine viral diarrhea virus 1 isolate MA_101_05 (GenBank No. LT968777.1). (Right) Alignment of amino acid sequences of NS3 of 37 Flaviviride virus. The four conserved amino acids are highlighted in yellow and the arginine is highlighted with blue spot.

S4 Fig. **qRT-PCR analysis of Dicer1, AGO1, AGO2, Tmr1, Tsn and Gemin4.** Analysis of major components gene expression of Dicer1, AGO1, AGO2, Tmr1, Tsn and Gemin4 using qRT-PCR in NAs infected with the indicated P3 after 48 hours.

S5 Fig. **Predication of RNA-binding sites in NS3 amino acid sequence using aaRNA.** The RNA binding sites on NS3 of JEV was analyzed using a PDB format of NS3 (PDB entry 2Z83) with aaRNA (https://sysimm.ifrec.osaka-u.ac.jp/aarna/), 4 arginine (R202, R226, R388 or R464 respectively) on the NS3 has a higher score (> 0.5) than other amino acid sites.

S6 Fig. **Predication of RNA-binding sites in NS3 amino acid sequence using Pprint.** The R202 and R464 that located in the helicase region of NS3 has high RNA interacting ability, which was analyzed by aaRNA (http://www.imtech.res.in/raghava/pprint/).

S7 Fig. **Predication of RNA-binding sites in NS3 amino acid sequence and miRNA nucleotide sequences using PRIdictor.** The R202 and R226 that located in the helicase region of NS3 has high RNA binding ability, which was analyzed by PRIdictor (http://bclab.inha.ac.kr/pridictor/).

S1 Table. **The proteins of NA cells identified by LC-MS from the co-immunoprecipitation with NS3.**

S2 Table. **Primers used for quantification of IL-1β and components of RISC mRNA.**

S3 Table. **The reverse transcription primers used for reverse transcribed all mature miRNA that were added with specific stem-loop structure at the end of 5’.**

S4 Table. **Primers used for quantification of pri-miRNA and pre-miRNA.**

S5 Table. **Primers used for quantification of mature miRNA and U6.**

